# Coordination between cytoskeletal organization, cell contraction and extracellular matrix development, is depended on LOX for aneurysm prevention

**DOI:** 10.1101/2024.02.23.581837

**Authors:** Rohtem Aviram, Shelly Zaffryar-Eilot, Anna Kaganovsky, Anas Odeh, Shay Melamed, Ruslana Militsin, Cameron B. Pinnock, Ariel Shemesh, Raz Palty, Santhi K. Ganesh, Peleg Hasson

## Abstract

Distinct, seemingly independent, cellular pathways affecting intracellular machineries or extracellular matrix (ECM) deposition and organization, have been implicated in aneurysm formation. One of the key genes associated with the pathology in both humans and mice is Lysyl oxidase (LOX), a secreted ECM-modifying enzyme, highly expressed in medial vascular smooth muscle cells. To dissect the mechanisms leading to aneurysm development, we conditionally deleted *Lox* in smooth muscle cells. We find that cytoskeletal organization is lost following *Lox* deletion. Cell culture assays and in vivo analyses demonstrate a cell-autonomous role for LOX affecting myosin light chain phosphorylation and cytoskeletal assembly resulting in irregular smooth muscle contraction. These results not only highlight new intracellular roles for LOX, but notably they link between multiple processes leading to aneurysm formation suggesting LOX coordinates ECM development, cytoskeletal organization and cell contraction required for media development and function.

## Introduction

Vascular smooth muscle cells (VSMCs) are the main cell type within the medial compartment of large vessels such as the aorta. These VSMCs are the main source for the secretion of extracellular matrix (ECM) and contraction, activities that are indispensable for vascular development, maintenance and regulation of blood pressure. Disruption of these VSMC activities results in thoracic aortic disease (TAD) manifesting in multiple vascular pathologies including aneurysms, dissections and stenosis.

Three main groups of pathogenic genetic variants affecting VSMCs have been associated with TAD ^1^. These include ECM structural proteins (primarily those composing or regulating the development of elastic fibers and collagens), members of the transforming growth factor β (TGFβ) signaling pathway and genes involved in VSMC contractile machinery ^2^. While an intimate link between structural components of the ECM and TGFβ signaling has been previously demonstrated ^3^, it is less clear why disruption of the VSMC contractile machinery results in aneurysm formation in a similar manner to that observed in connective tissue (or ECM) pathologies and what are the mechanisms that link between these causes.

LYSYL OXIDASE (LOX) is a key enzyme belonging to the lysyl oxidase family of secreted ECM modifiers consisting of LOX and LOX-Like (LOXL) 1-4. LOX is highly expressed in VSMCs and is the predominant family member in the aortic media ^4,5^. LOX is secreted as a proenzyme and is cleaved by various proteases. Following cleavage, both the mature active enzyme and its N-terminal propeptide (LOX-PP) are released and mediate distinct functions. Previous work has demonstrated that in VSMC both LOX-PP and the mature active enzyme can enter the cell and exert various activities ^6–9^. Key primary targets for LOX enzymatic activity within the media are elastin and collagens where it functions as the main enzyme mediating their crosslinking. Accordingly, mutations affecting LOX expression or enzymatic activity lead to disruption of ECM integrity and are associated with aneurysms in both humans and mice ^10–13^. Surprisingly, although LOX is one of the most prominent genes linked with TAD, no direct genetic assessment of its role specifically in VSMC has been carried out in vivo.

To dissect mechanisms that underlie aneurysm development and involvement of VSMC in the process, we used a conditional knockout strategy to delete *Lox* using Cre lines expressed under the *Myh11* promoter, a promoter specifically expressed in SMCs. Using this strategy and knockdown assays in human aortic SMCs (HAOSMC) we find that LOX plays a key cell-autonomous role in regulating VSMC cytoskeleton in an ECM-independent manner. Disruption of this latter activity results in deregulation of SMC contraction in the aortic media. Our results highlight a hitherto unappreciated role for LOX and suggest that apart from the observed ECM defects, disruption of its intracellular activities lead to abnormal cytoskeletal organization, deregulated contraction, and inability to respond to altered ECM. Surprisingly, we find the latter activities are enzyme- and ECM-independent. Our results therefore highlight a missing link between the three distinct gene groups associated with aneurysms, thus serving as a molecular paradigm for the development of phenotypes that culminate in aneurysm.

## Results

### VSMC-specific Lox deletion leads to aneurysms

To dissect the mechanisms promoting TAD that could result in aneurysms, we set to specifically delete *Lox*, an ECM modifying enzyme highly associated with aneurysms, in VSMC using a conditional knockout strategy. Towards that end, we crossed the *Myh11:Cre^ERT2^*deleter line ^14^ to mice bearing a *Lox* conditional knockout allele (*Lox^fl^*) ^15^. Upon weaning, at the age of 3 weeks, *Myh11:Cre^ERT2^; Lox^fl/fl^* mice (or *Lox^fl/fl^*as controls) were IP injected with tamoxifen (100µl, 20mg/ml) for 5 consecutive days and then on once a month. Surprisingly, we find that even though aortic Lox levels were significantly downregulated (~65-90% deletion) as monitored by a western blot (WB) analysis (Supp. Fig.1A,B), no gross observable phenotypes were detected in the aortas of the mutant mice, no changes to their blood pressure was noted, and their lifespan was, similarly, not affected.

The surprising lack of visible phenotype following *Lox* deletion in the VSMC led us to test whether such aortas lacking Lox activity in their media could withstand hypertension. Towards that end, at the age of 3 months, osmotic pumps (alzet) infused with Angiotensin II (AngII; 10mg/kg/day) were subcutaneously implanted for the duration of one month to increase blood pressure. Blood pressure was monitored after 8 days and in all cases no differences between control (*Lox^fl/fl^*) and mutant (*Myh11:Cre^ERT2^; Lox^fl/fl^*) mice were observed (Fig. 1A) altogether demonstrating that the loss of Lox from the VSMC did not affect their ability to respond to AngII and become hypertensive. Following one month, at 4 months of age, mice were harvested and their aortas dissected. WB analysis demonstrates AngII infusion did not affect the extent of Lox deletion (Fig. 1B). While among the AngII-treated control mice (n=9; *Lox^fl/fl^*), only one displayed an aneurysm, all mice with reduced medial Lox [of the genotypes *Myh11:Cre^ERT2^; Lox^fl/fl^* or *Myh11:Cre^ERT2^; Lox^fl/Δ^* (n=10)], displayed at least one aneurysm (Fig. 1C). Aneurysms were located throughout the aorta, in the abdominal regions as well as in the thoracic and arch regions. Altogether, these results demonstrate that Lox activity in the media is essential for the withstanding of hypertensive conditions.

**Figure 1.**
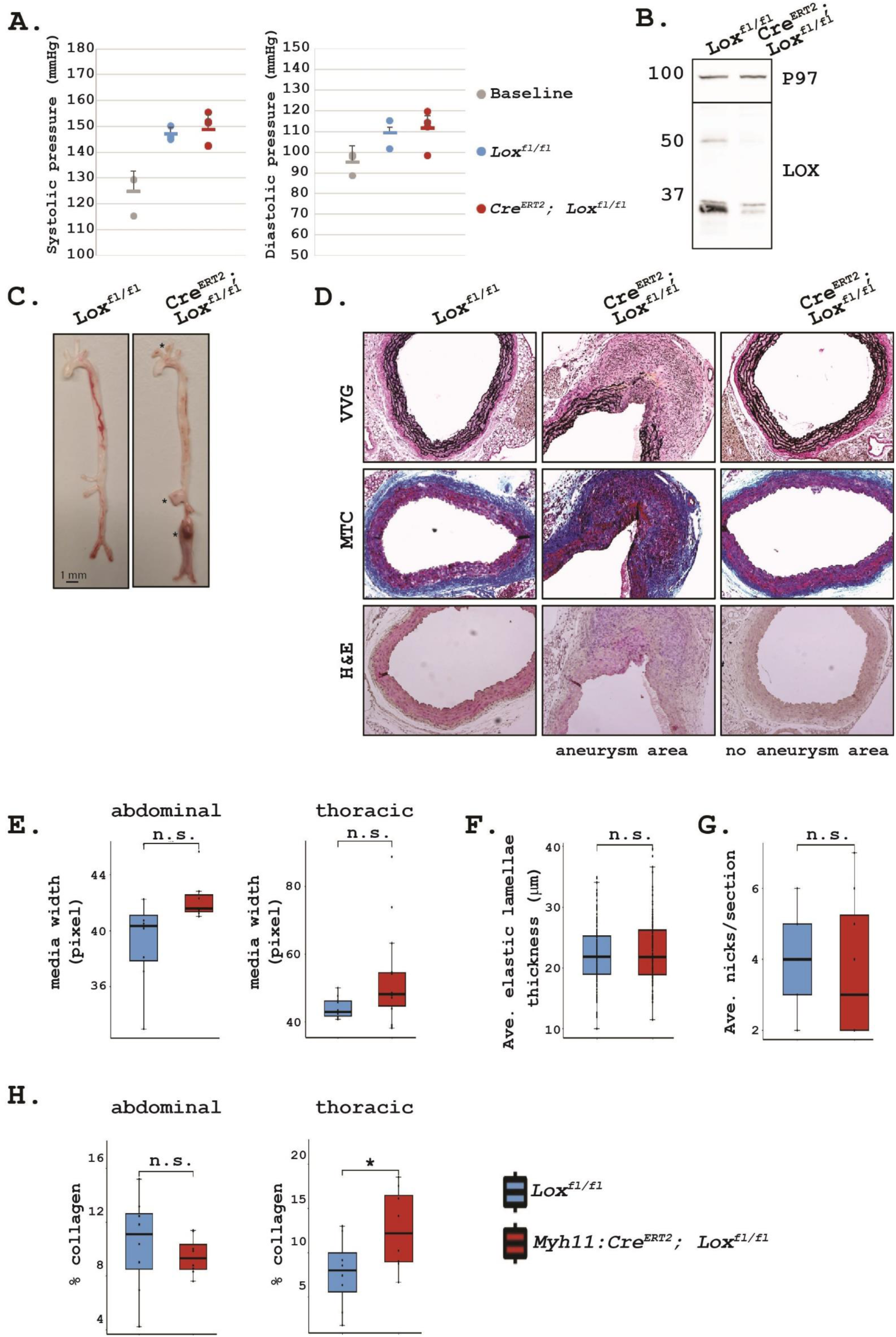
*Lox* knockout in VSMC promotes aneurysm formation. Blood pressure measurements of control (*Lox^fl/fl^*) prior to and following AngII treatment as well as of *Myh11:Cre^ERT2^;Lox^fl/fl^*mice (A). Western blot for Lox of control and *Myh11:Cre^ERT2^;Lox^fl/fl^*mouse aortas following AngII treatment (B). Aortas of control (left) or *Myh11:Cre^ERT2^; Lox^Δ/fl^*(right) infused with AngII (C). Note aneurysms in *Myh11:Cre^ERT2^; Lox^Δ/fl^* (marked with *). Histological staining of *control* (left) and *Myh11:Cre^ERT2^;Lox^fl/fl^*(middle and right) aortic sections for Verhoeff Van-Gieson (VVG), Mason’s Trichrome (MTC) and H&E. Note in control aortas, properly formed elastic fibers and defined tissue layers in contrast to the area of aneurysm in mutant mice, with torn elastic fibers and cell infiltration at point of rupture (middle column). In contrast, in non-aneurysmal areas of mutant aorta (right) no defect in media width (E), elastic lamella thickness (F) or nicks within (G) were observed. Collagen deposition was not affected in abdominal regions yet was upregulated in thoracic ones (H) in comparison to control.

Histological examination of the aneurysms, regardless of their location, demonstrated that as expected, elastic lamellae marked by Verhoeff-Van Gieson (VVG) staining were severely affected highlighting multiple tears; Masson’s Trichrome (MTC) staining revealed increased collagen deposition and infiltration of immune cells to the diseased region (Fig. 1D). Interestingly, upon monitoring non-aneurysmal regions, no significant differences were observed in media width, average thickness of elastic lamella and number of breaks within them, although collagen deposition was significantly increased in thoracic regions of the mutant aorta irrespective of aneurysm location (Fig. 1D-H). These observations were surprising, since *Lox* was depleted throughout the aortic media, raising the hypothesis that although its deletion affected the VSMC-dependent ECM and elastin crosslinking, it may have resulted in additional, possibly ECM-independent processes that contributed to the aneurysms.

### LOX regulates multiple cellular behaviors

To identify additional roles for Lox in VSMC, we took advantage of human aortic smooth muscle cells (HAOSMC). These primary cells express high LOX levels and in a similar manner to that observed in mice ^4,5^, it serves as the predominant family member in these cells (Fig. 2A). HAOSMC secrete ECM ^16^ (see below) and can be easily manipulated in culture. To test if LOX levels can be knocked-down we infected the cells with lentiviral particles expressing small hairpin sequences targeting *LOX* (*shLOX*) or as control, a sequence with no target gene (*shCtrl*). We find that following *LOX* knockdown, LOX protein levels were significantly downregulated without considerably affecting the other members of the family (Fig. 2B,C). We also infected the cells with two more *shLOX* lentiviral independent particles (marked as s*hLOX2* and *shLOX3*) where similar inhibition of LOX expression was observed (Fig. 2C). Reduction in LOX expression via *shLOX* resulted in a significant decrease in cell size (Fig. 2D-F). A comparable reduction in HAOSMC cell size following *shLOX2* and *shLOX3* transfection was also observed (Fig. 2F). Thus, suggesting the decrease in cell size phenotype was specific to *LOX* and not resulting from a non-specific target inhibited by the small hairpin sequences.

**Figure 2.**
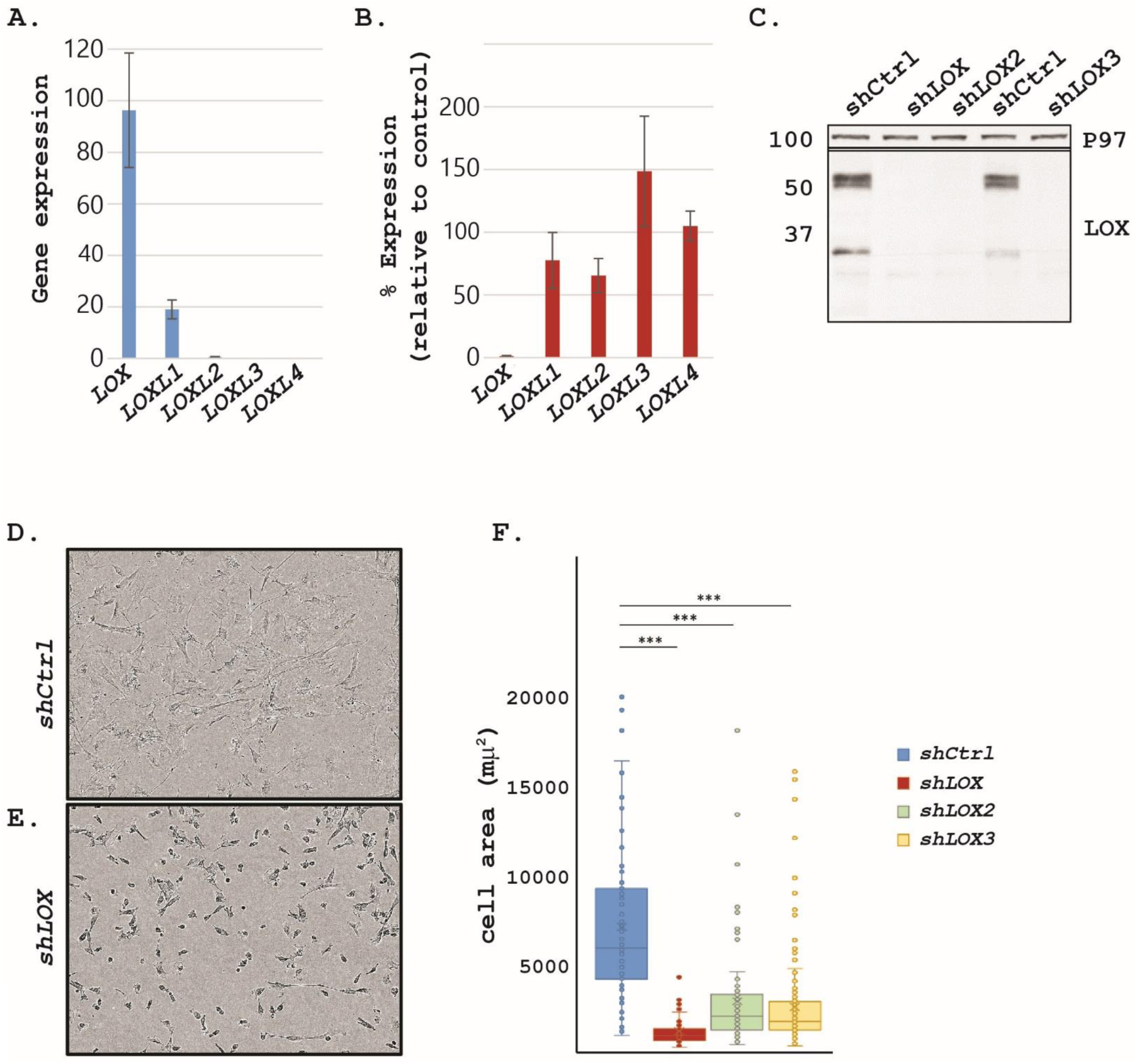
*Lox* knockdown in HAOSMCs results in smaller cells. Quantitative RT-PCR for *LOX* family members shows *LOX* is the predominant member of the family in HAOSMCs (A) Quantitative RT-PCR for *LOX* family members demonstrate they are not significantly affected by *LOX* deletion in HAOSMC (B). Western blot for LOX following knockdown with three distinct shRNA constructs demonstrate they all lead to significant LOX knockdown (C). Bright-field images of *shCtrl* (D) or *shLOX* (E) cells shows HAOSMCs devoid of *LOX* are smaller. Quantification of cell area reveals significant reduction in the 3 *shLOX* sequences (F).

The association of LOX with aneurysms has been suggested to be dependent on its enzymatic activity where it regulates medial ECM formation and stabilization ^5,10,12,17^. We therefore set to test whether the observed HAOSMC phenotypes are likewise enzyme-dependent by adding increasing concentrations of up to 500 µM of the pan-LOX inhibitor βAPN ^18^ or 10 µM of the LOX-specific CCT365623 inhibitor ^19^. Neither of these treatments affected LOX protein levels and surprisingly they had no effect on HAOSMC cell size (Supp. Fig. 2A,B).

The lack of LOX enzymatic involvement raised the possibility that the observed phenotype might be an activity associated with the LOX N-terminal region, LOX-PP, previously demonstrated to inhibit VSMC proliferation ^9^. However, in contrast to the phenotypes previously related to LOX-PP, immunostaining for the proliferation associated marker Ki-67 demonstrated HAOSMC depleted of LOX proliferated poorly (Supp. Fig. 2C,D). Live cell imaging over a 3-day period not only reinforced the poor proliferation of the LOX-devoid cells, but also demonstrated they exhibited increased cell death (Supp. Fig. 2E,F) altogether suggesting the observed phenotypes in the *LOX* depleted cells are distinct from those associated with LOX-PP. Interestingly, while the majority of nuclei in *shCtrl* cells appeared to be in a disc-like shape where only ~14% had an irregular contour, 50% of the nuclei of *shLOX* cells were irregular (Supp. Fig. 2G,H). Overall, these results suggest *LOX* depletion affected a general cellular property/properties independent of its enzymatic activities.

### Autonomous LOX phenotypes

LOX is a key ECM-modifier enzyme, hence we wished to test whether its depletion also affected the ECM the cells secrete. Towards that end, we cultured the *shCtrl* and *shLOX* HAOSMC for 8 days, removed the cells, and the ECM they have secreted was then submitted to LC-MS/MS analysis. As expected, significant differences were observed in ECM protein levels between the control and LOX-depleted cells (Supp. Fig. 3A). Having seen these differences, we wished to test whether the observed phenotypes could be attributed to the abnormal ECM secreted by the LOX-depleted cells by assaying whether proper ECM could rescue them. Therefore, control HAOSMC were cultured for 8-10 days and allowed to form a tight surface of compact cells and ECM (Supp. Fig. 3B). Following culturing, cells were removed without affecting the ECM they have secreted which remained bound to the tissue culture plates (Supp. Fig. 3C). *shCtrl* or *shLOX* HAOSMC were then seeded directly onto the ECM (or onto standard tissue culture plates as control) and cultured for 24 hours. Following plating *shCtrl* cells directly on plastic, the cells spread yet they do not assume any directionality (Fig. 3A). In contrast, once they are plated onto mature ECM, they rapidly elongate and assume the orientation of the ECM, i.e. that of the cells that have previously laid it (Fig. 3B,C). In contrast, *shLOX* cells cultured on similarly mature ECM were comparable in size and shape to those cultured directly on plastic (Fig. 3A,B). Further these LOX-devoid cells did not elongate neither did they assume the orientation of the ECM they were laid upon (Fig. 3B,C). Altogether, these results suggest the observed LOX-dependent phenotypes are not caused due to an abnormal ECM laid by the mutant cells.

**Figure 3.**
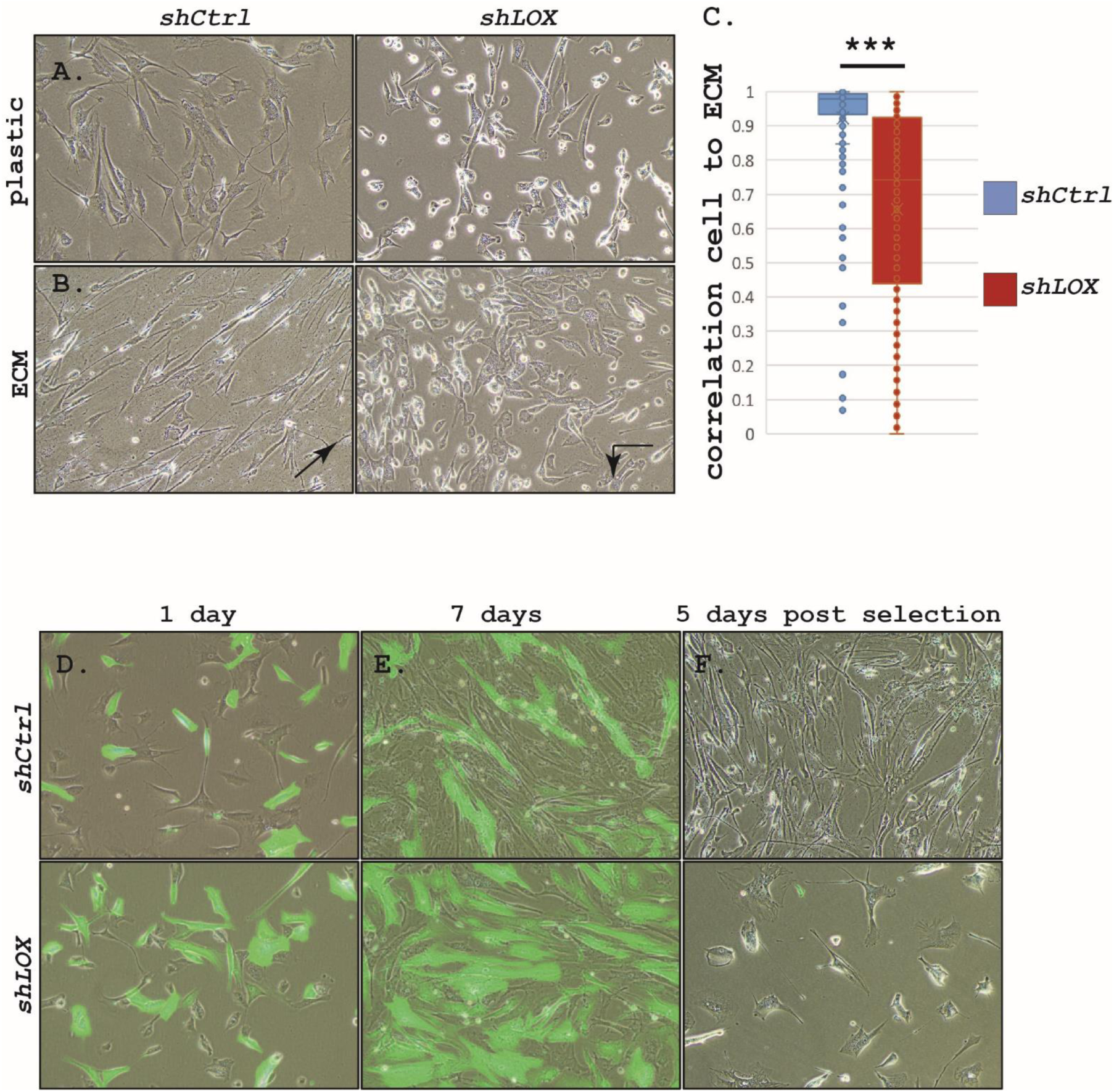
LOX is autonomously required in HAOSMCs for interactions with the ECM. *shCtrl* (left) or *shLOX* (right) cells cultured directly on plastic (A) or on HAOSMC-derived ECM (B). While *shCtrl* assume directionality of ECM they lay on (arrow in B, left), that of *shLOX* cells does not correlate to the ECM (arrow in B, right). Quantification of cell-to-matrix correlation shows significant differences between the cell genotypes (C). Bright-field and fluorescent images of GFP-tagged HAOSMCs co-cultured with *shCtrl* cells (D, top) or *shLOX* cells (D, bottom) and monitored for a week (E top and bottom, respectively). After 7 days the GFP cells were killed using puromycin selection leaving culture dishes with *shCtrl* (F, top) or *shLOX* (F, bottom) cells.

While LOX is mostly considered as a secreted enzyme where many of its activities are carried out within the extracellular environment, previous work has suggested that cells, and primarily SMC, can uptake exogenous mature enzyme ^6^. We therefore reasoned that co-culturing LOX-depleted cells with ones that secrete LOX, would rescue, at least to some extent, the observed phenotypes (e.g., small cell size and reduced proliferation). Towards that end, GFP-transfected HAOSMC were plated at a concentration of 1:1 and cultured with *shLOX* or *shCtrl* cells for one week. Cells were imaged throughout this period. By the end of the week, the GFP-expressing HAOSMC cells occupied ~50% of the dish when cultured with *shCtrl* cells suggesting they proliferated at similar rates. In striking contrast, plates in which the *shLOX* and GFP-expressing HAOSMCs were co-cultured, occupied primarily the GFP-expressing cells. These results demonstrate that although the latter cells secreted LOX (Supp. Fig. 3D), the exogenous enzyme was not able to rescue the proliferation or spreading of the *LOX*-depleted cells (Fig. 3D,E; Supp. Fig. 3E). To test whether the reduced cell size of the *shLOX* cells was not caused by their inability to compete with the otherwise wild-type GFP-expressing cells, we eliminated the latter GFP-marked cells as they did not express any resistance marker. Following the addition of puromycin, only the *shCtrl* or *shLOX* cells which stably express the knockdown construct survived. We find that following the removal of the GFP-expressing cells, *shCtrl* cells spread, elongated and occupied the majority of the culture. In contrast, the *shLOX* cells hardly proliferated and maintained their small size (Fig. 3F). Overall, the combined results suggest the observed LOX-dependent phenotypes, which cannot be compensated by exogenous LOX, are cell autonomous, and regulate the cells’ ability to sense and respond to distinct ECM environments.

### LOX associates with cytoskeletal elements

To test what could be the mechanism underlying the observed small cell size, defective nuclear shape and the inability to respond to an ECM we set to identify LOX partners in the VSMC. We immunoprecipitated LOX from HAOSMC cell lysates and submitted the precipitate to a LC-MS/MS analysis (Supp. Table 1). Proteins identified by at least 3 distinct peptides, enriched at least 4 fold with a p-value <0.05 in comparison to the control IgG precipitate were then submitted for bioinformatics analysis. Kyoto Encyclopedia of Genes and Genomes (KEGG) analysis identified significant enrichment of cytoskeleton-associated proteins to be precipitated with LOX (Supp. Fig. 4A). Monitoring cellular functions using Ingenuity Pathway Analysis (IPA) further reinforced the significant enrichment of cytoskeletal elements among the proteins complexed, directly or indirectly, with LOX (Supp. Fig. 4B).

The significant enrichment of cytoskeletal proteins associated with LOX containing complexes, prompted us to monitor cytoskeletal arrangement in the *shLOX* HAOSMC. Staining for phalloidin marking F-actin demonstrates significant disruption of actin organization. Although stress fibers are present in the *shLOX* HAOSMC, many of the fibers are disrupted and do not extend the cells’ width or length as in the *shCtrl* HAOSMC (Fig. 4A). Immunostaining for tubulin reveals that in a like manner to F-actin, the microtubule network was also disrupted and disorganized (Fig. 4B). High-resolution transmission electron microscope (TEM) imaging further demonstrates that cytoskeletal bundles are significantly disrupted in the *shLOX* cells (Fig. 4C).

**Figure 4.**
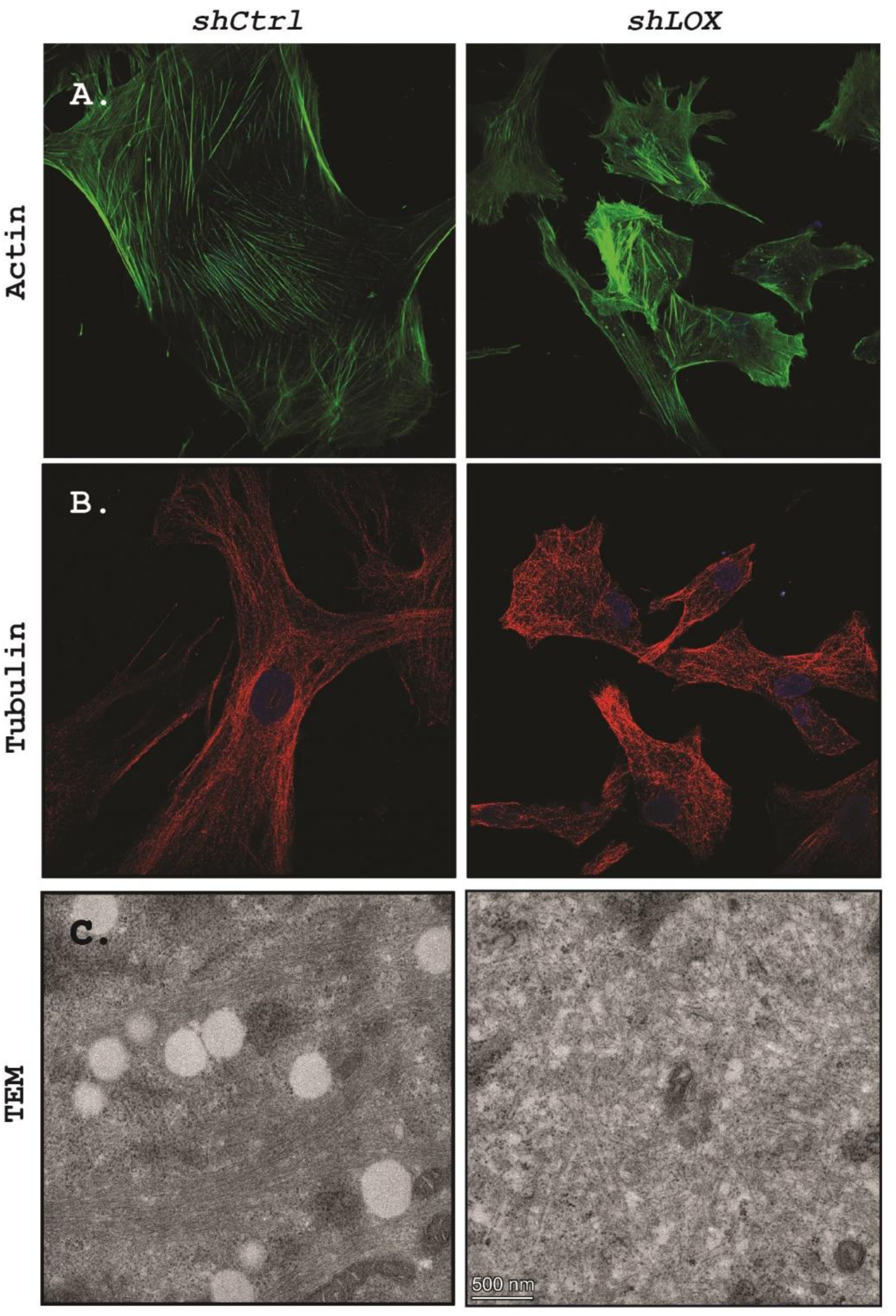
*LOX* knockdown in HAOSMCs affects cytoskeletal organization. Staining for actin (phalloidin, green; A) and tubulin (red; B) in *shCtrl* (left) and *shLOX* (right) HAOSMC. High magnification TEM images of *shCtrl* (left) and *shLOX* (right) shows cytoskeletal organization is significantly affected following *LOX* knockdown (C).

To further dissect the disruption to the cytoskeleton, we monitored actin-binding proteins that are abundant in VSMC. Smoothelin (SMTN), an actin binding protein aligns with the stress fibers in control cells. However, following *LOX* knockdown this pattern is lost and SMTN is found scattered within the cytoplasm (Fig. 5A). Notably, this disruption is not specific to SMTN; monitoring other actin-binding proteins namely MyosinIIB, Calponin, SM22 (Transgelin; TAGLN) and Filamin A, all revealed a similar pattern; while in the *shCtrl* cells these proteins aligned with the stress fibers, in the *shLOX* cells, expression of these proteins was dispersed throughout the cytoplasm and barely aligned with the fibers (Fig. 5B-E). Quantification of Smoothelin and SM22 expression demonstrated they were not significantly affected by the loss of LOX (Fig. 5F-H and LC-MS/MS analysis, Supp. Table 2), indicating that cytoskeletal organization, rather the expression levels of its components, are affected by the loss of LOX.

**Figure 5.**
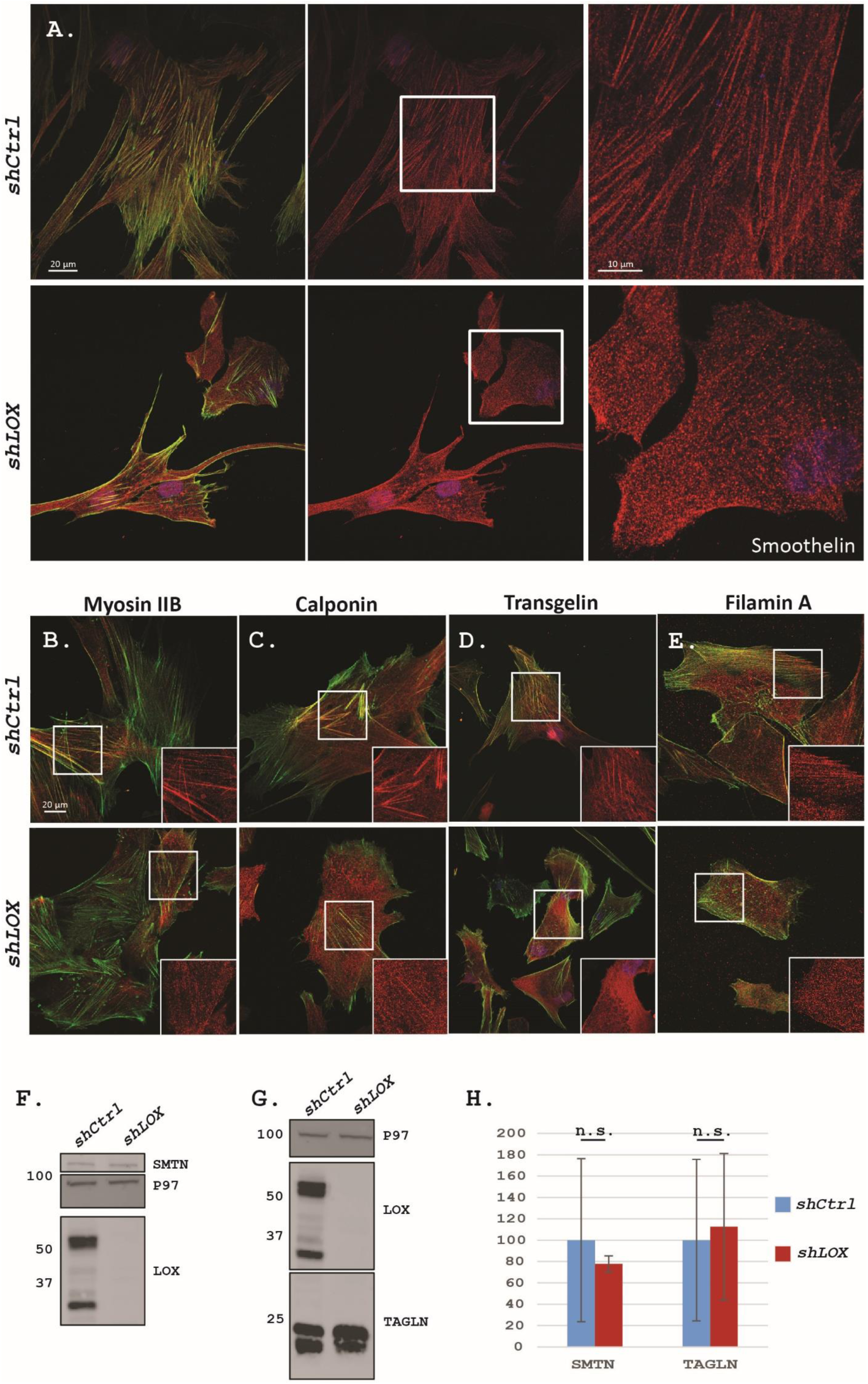
Localization of actin binding proteins is disrupted following LOX knockdown. Confocal images of *shCtrl* (A, top) and *shLOX* (A, bottom) HAOSMCs immunostained for for Smoothelin (red) and phalloidin (actin, green). Note Smoothelin localization to actin fibrils is lost in *shLOX* cells (higher magnification of boxed area in right panel). Confocal images of Myosin IIB (B), Calponin (C), Transgelin (D) and Filamin A (E) in *shCtrl* (top) and *shLOX* (bottom) HAOSMC shows similar disruption of actin binding proteins in cells devoid of LOX. Western blots for Smoothelin (F) and Transgelin (G) and corresponding quantification (H) in *shCtrl* and *shLOX* cells shows their levels are not affected following *LOX* knockdown.

### MLC phosphorylation is abnormally regulated in LOX mutant cells

The small, gaunt appearance of the *shLOX* cells along with their abnormal cytoskeletal organization raised the possibility that contraction of their actin cytoskeleton might be abnormally regulated. The regulatory myosin light chain (MLC/Myl9) is the main regulatory protein controlling actin contraction and phosphorylated MLC (p-MLC) is found on the cytoskeletal contractile units. MLC is activated by phosphorylation that in turn is dually regulated by a phosphatase (MLCP) and a kinase (MLCK)^20^. Deregulated MLC phosphorylation has been associated with aneurysms and cardiovascular pathologies ^20,21^. MLCP is inhibited by the Rho associated kinases ROCK1 and ROCK2 hence these two kinases serve as positive regulators of filamentous actin buildup. In addition, they have been shown to directly phosphorylate actin binding proteins regulating their association with F-actin ^22–24^ altogether raising the hypothesis they may be involved in the above phenotypes. LC-MS/MS analysis and WB of *shCtrl* and *shLOX* HAOSMC demonstrated both these kinases are significantly down regulated in the latter (Supp. Table 2; Fig. 6A,B). Culturing *shCtrl* and *shLOX* cells with a ROCK inhibitor mostly affected the *shCtrl* cells but not the *shLOX* ones in agreement with the latter having an already reduced ROCK1/2 expression (Fig. 6C,D). As expected, ROCK inhibition in *shCtrl* cells significantly disrupted their F-actin and in the filaments that were still present, no Smoothelin alignment was observed, suggesting this is a ROCK-dependent process (Fig. 6D). Accordingly, the culturing of the cells with a Rho activator II, also had little effect on the *shLOX* cells since ROCK proteins, the downstream effectors of this activator, were not present for it to properly function (Fig. 6E).

**Figure 6.**
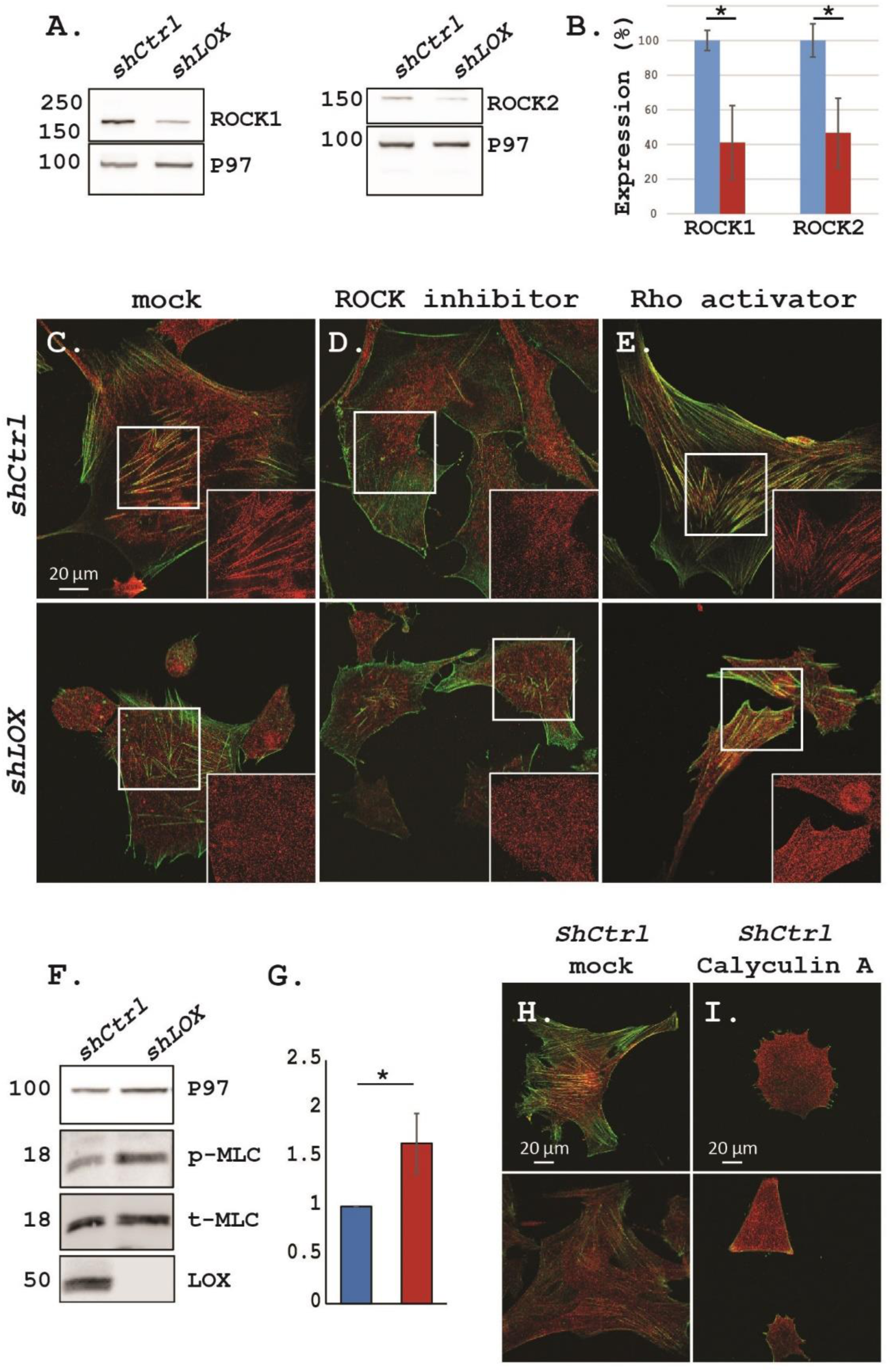
*LOX* knockdown affects the Rho/ROCK pathway. Western blot (A) and quantification (B) of ROCK1 and ROCK2 in *shCtrl* and *shLOX* HAOSMC shows ROCK proteins are downregulated following LOX deletion. Confocal images of immunostained HAOSMC for Smoothelin (red) and phalloidin (actin, green). Mock (C), ROCK inhibitor (D) or Rho activator (E) treated *shCtrl* (top) or *shLOX* (bottom) cells. Smoothelin localization to actin filaments is lost in *shCtrl* cells following ROCK inhibition (D) and is not regained following Rho activation in *shLOX* cells (E). WB (F) and quantification (G) of total and p-MLC in *control* and *shLOX* cells. Confocal images (2 images each, phaloidin green; Smoothelin, red) of untreated (H) and Calyculin A treated (I) *shCtrl* HOASMC demonstrate inhibition of MLC-phosphatase leads to reduced cell size and cytoskeletal defects.

The observation that the expression of the ROCK proteins was reduced in the LOX-devoid cells suggested MLC activation, i.e. its phosphorylation, should be deregulated. WB for phosphorylated MLC demonstrates upregulated phosphorylation (Fig. 6F,G). Culturing the *shCtrl* HAOSMC with Calyculin A, a MLCP inhibitor, resulted in small *shLOX*-like cells with few organized F-actin filaments (Fig. 6H,I) but also led to a significant increase in *shLOX* cell death (not shown). Altogether, these results suggest that MLC activation is abnormally regulated in a ROCK/MLCP-independent manner in the *LOX*-depleted cells.

We therefore set to test whether the other MLC regulator MLCK, a calcium-dependent kinase, could be aberrantly regulated in the *shLOX* cells. MLCK is activated following calcium influx into the cytoplasm from the cellular storages (e.g., sarcoplasmic reticulum) or from the extracellular space, hence we monitored calcium homeostasis in the *LOX* knockdown and control cells. Culturing the HAOSMC cells in calcium-free Ringer’s solution and 0.2mM EGTA (a Ca^2+^ chelator), allowed us to monitor calcium cellular homeostasis in the *shCtrl* and *shLOX* cells. Using Fura-2 AM, a fluorescent Ca^2+^ indicator and ionomycin, a Ca^2+^ ionophore that induces release of stored Ca^2+^ to the cytoplasm, we traced the levels of cytosolic calcium before and after release of stored Ca^2+^. We find that *LOX* knockdown does not change resting Ca^2+^ levels. However, while calcium levels in cellular stores is also not affected (Fig. 7A), tracing the calcium levels demonstrates that the ability of the *shLOX* cells to restore cytosolic calcium back to resting levels as monitored by the rate of return to a steady state, is significantly inhibited (Fig. 7B-E). This finding suggest that following a variety of physiologically relevant stimuli, *shLOX* treated cells would experience prolonged periods of elevated cytosolic calcium and consequently abnormal MLCK activation. Should MLCK activation be one of the causes of the observed phenotypes, we expect that its inhibition will improve, at least to some extent, the *shLOX* phenotypes. Indeed, culturing the cells (*shCtrl* or *shLOX* HAOSMC) for four days with a MLCK inhibitor resulted in a cell size increase and the re-appearance of stress fibers in *shLOX* cells. In contrast, this treatment slightly worsened the appearance of the control cells (Fig. 7F-H) altogether reinforcing the notion that MLCK is abnormally regulated in the *shLOX* HAOSMC.

**Figure 7.**
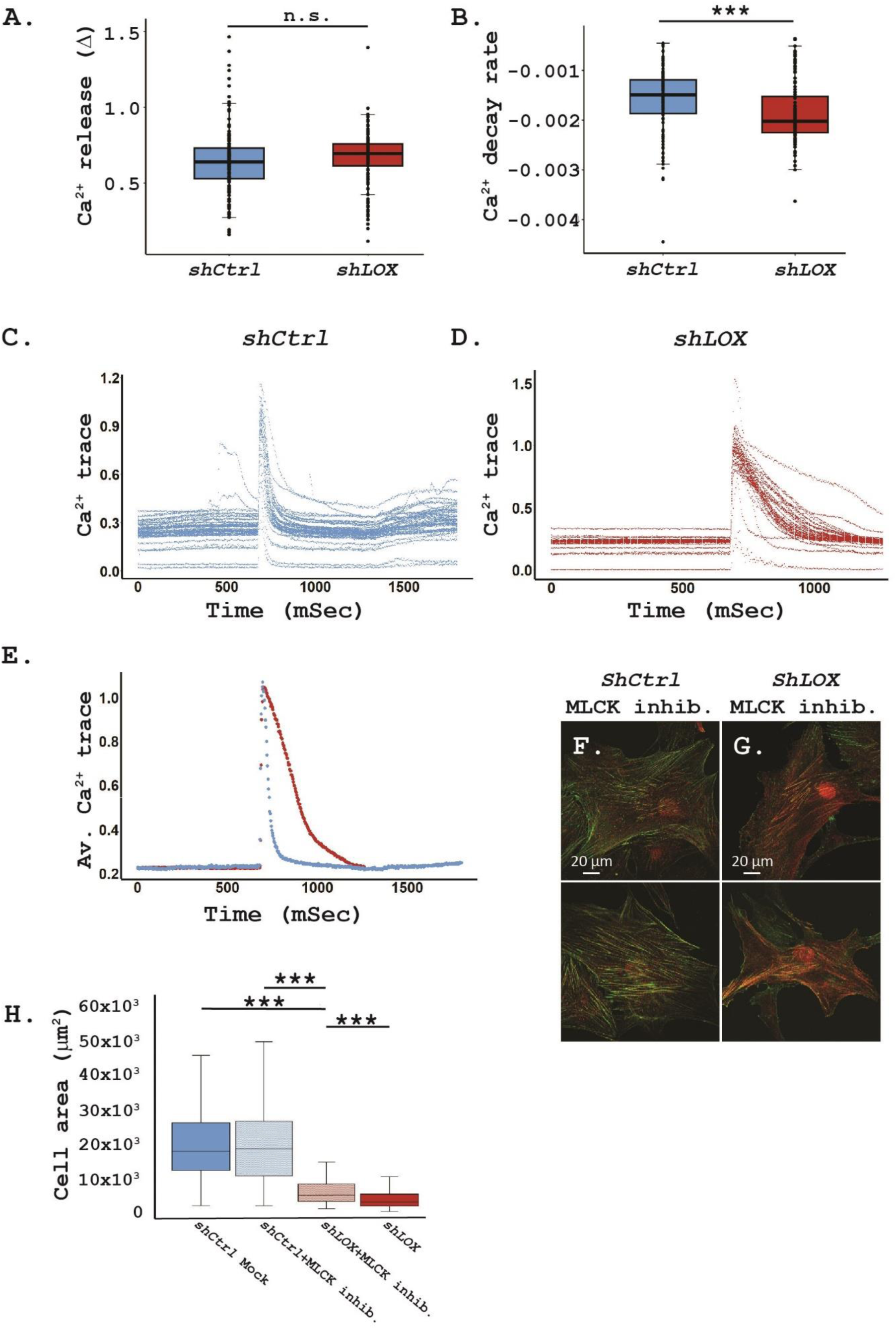
LOX-deleted cells exhibit disrupted calcium homeostasis. Intracellular calcium tracing show rate and magnitude of calcium release from intracellular storages and extracellular media are not altered in *LOX*-depleted cells (A). In contrast, the rate of calcium re-entry back into storages or its extracellular secretion is significantly inhibited in mutant cells (B). Single cell readings from *shCtrl* (C) and *shLOX* (D) as well as corresponding averages (E) demonstrate reduced rate of cytoplasmic calcium removal. Confocal images for phaloidin (green) and Smoothelin (red) of *shCtrl* cells with MLCK inhibitor results in minor defects of actin filament linearity (F, 2 images), whereas partially rescues *shLOX* cytoskeletal organization (G, 2 images) and leads to an increase in their cell size (H).

The above observations were all carried out in cultured cells. Therefore, we next set to examine whether these observations are relevant also to *Lox* roles in vivo in SMCs. We took advantage of *Myh11:Cre^EGFP^* ^25^ which enables the deletion of a floxed gene in SMC from the onset of *Myh11* expression without any requirement for Tamoxifen induction and monitored whether the cytoskeleton is affected following this *Lox*-deletion regime. As with the Tamoxifen-induced Cre, aneurysms were observed only following AngII infusion in *Myh11:Cre^EGFP^; Lox^fl/fl^* mice (n=5, Supp. Fig. 5) while non-hypertensive mice were viable, normally behaved and did not show any overall phenotype demonstrating *Lox* deletion in SMC on its own even from early stages is not sufficient for aneurysm formation. However, high resolution TEM imaging of medial SMCs demonstrated cytoskeletal elements were significantly disorganized in the mutant aortas even without AngII-mediated hypertension (Fig. 8A-C, higher magnification in A’-C’). Interestingly, upon monitoring p-MLC immunostaining on aortic sections, we find that in control mice p-MLC immunostaining intensities are rather uniform throughout the aortic media. In contrast, although absolute values of p-MLC reach higher levels in *Myh11:Cre^ERT2^; Lox^fl/fl^*, immunostaining reveals a more variable pattern of p-MLC expression (Fig. 8D-F). Spatial quantification of immunostaining intensities show higher differences between the median and mean values in the mutant aortas (|median-mean|=3.69 in control vs. 7.78 in mutant aortas; n=4 aortas of each genotype) (Fig. 8F) demonstrating p-MLC inconsistent activation. Since p-MLC serves as a readout of cell contraction, these observations suggest that SMC contraction in the *Lox* mutant aortas is scattered and irregular.

**Figure 8.**
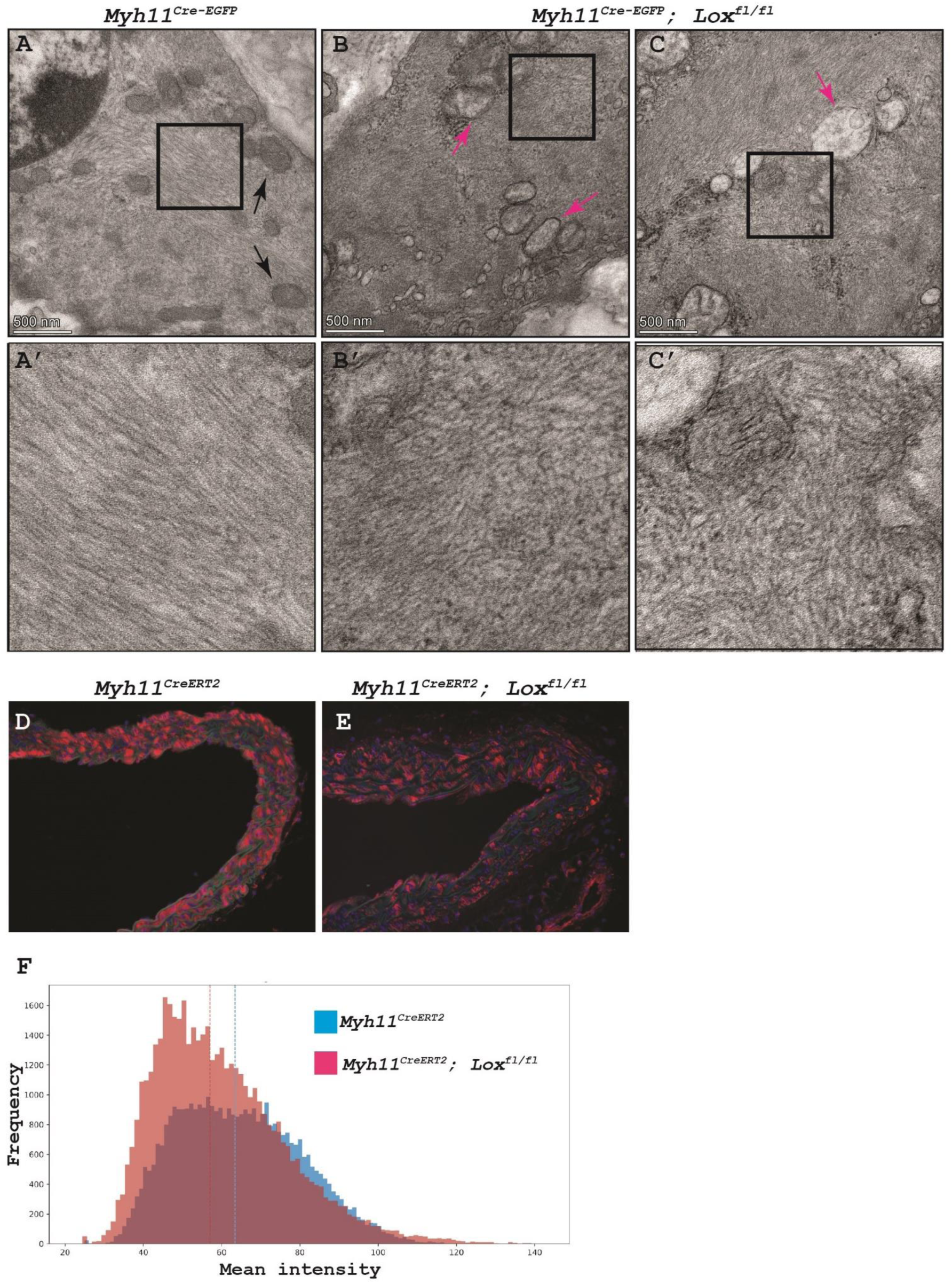
Lox regulates cytoskeletal organization in vivo independently of hypertension. High resolution TEM images of aorta sections (A, control; B, C *Myh11:Cre^EGFP^; Lox^fl/fl^*) demonstrate lack of cytoskeletal organization in *Lox* knockout SMC even without hypertension. Black and pink arrows point to healthy and abnormal appearing mitochondria, respectively. Higher magnification of boxed areas A’-C’. Immunostaining for p-MLC (red) in control (D) and mutant (E) aortas demonstrates scattered and irregular cell activation. Histogram of p-MLC mean intensity frequency (n=4 of each genotype) demonstrate higher scattering of staining in *Myh11:Cre^ERT2^; Lox^fl/fl^* mutant aortas than in the control (*Myh11:Cre^ERT2^*) (F). Dotted lines indicate median values.

Apart from regulating cell contraction, cytosolic Ca^2+^ also regulates multiple other cellular processes and its excessive levels are associated with increased mitochondrial deformities ^26,27^. TEM imaging monitoring control and mutant aortas (n=4 mice of each genotype) demonstrate that while in control mice ~4% of the mitochondria had a damaged appearance (n=93), in *Myh11:Cre-EGFP; Lox^fl/fl^* >25% were damaged (n=182; Fig. 8A-C, black and pink arrows; Supp. Fig. 6). These results suggest that the observed scattered p-MLC activation might be due to reduced mitochondrial activity stemming from the increased cytoplasmic Ca^2+^.

## Discussion

Regulation of blood vessel integrity is key for maintaining proper vascular function. Disruption of extracellular elements such as the elastic fibers were found to play a major role in the emergence of weak spots, susceptible to rupture. Much work has focused on how cells within the media are affected by defects in the ECM, however aberrant intracellular processes that can in turn affect ECM and result in TAD have been given less attention. Administration of *β*APN leading to a pan lysyl oxidase enzymatic inhibition, *LOX* mutations in humans and *Lox* knockout mice demonstrated this enzyme, known for its role in elastin crosslinking, plays essential roles in the vasculature and its inhibition or loss results in ECM defects that promote TAD ^10,12,13^. Using a conditional knockout strategy, we were able to directly examine Lox roles in VSMCs in vivo. By focusing on the regions where no aneurysms occurred combined with cell culture assays, we demonstrate that LOX plays an additional, as yet unappreciated, major intracellular role in regulating VSMCs cytoskeleton affecting their contractile machinery.

Our finding that, intracellularly, LOX is highly enriched with cytoskeletal complexes along with the significant cytoskeletal abnormalities observed following its loss prompted us to further explore its involvement in the process. Phosphorylation of the myosin light chain serves as a key regulatory event in the formation of the contractile units of actin cytoskeleton and as such is tightly regulated via two main branches – ROCK1 and ROCK2 regulating MLC phosphatase on the one hand and a calcium-dependent kinase on the other. Surprisingly, although we find that in response to *LOX* knockdown ROCK1 and ROCK2 expression are significantly reduced, a process which could lead to enhanced phosphatase activity, ectopic MLC phosphorylation was observed raising the hypothesis that the calcium-dependent MLCK could be abnormally regulated. Probing calcium homeostasis in *shLOX* cells demonstrated significant delays in the ability of these cells to remove cytoplasmic calcium, suggesting it could underlie the observed prolonged MLCK activity in HAOSMC devoid of LOX. However, in vivo, rather than an increase in MLC phosphorylation, irregular and more scattered activation was observed. While this difference could be due to the higher complexity in vivo with multiple feedback loops, our finding that mitochondrial damage was more apparent in the *Lox*-mutant SMC could explain this observation.

Notably, hypertension induces vascular remodeling and SMC contraction ^28^, two processes which also involve LOX. Previous reports have further implicated LOX as a regulator of TGFβ signaling ^29–31^ but also as a downstream target for the pathway ^32^. Since TGFβ signaling is implicated in aneurysm formation ^2,33^ our observations, therefore, place LOX as a common player in the various processes implicated in aneurysms.

Our results further suggest that not only are LOX activities spatially separated, but they also differ in their dependence on its catalytic activities; while the intracellular regulation of cytoskeleton is mediated via non-catalytic activities, extracellular ECM-crosslinking is enzyme-dependent. Although LOX is an essential ECM-modifying enzyme, its intracellular roles are crucial for VSMC viability and for the integrity of the aortic tissue. This work therefore suggest that in response to reduced LOX expression, abnormal cytoskeletal organization leads to irregular smooth muscle cell contraction and this, along with mild ECM defects, even in the lack of hypertension, predispose the aorta to aneurysms. Thus, in agreement with previous findings where heterozygous *Lox* mice were viable and did not show any or only a mild phenotype ^5,10^ it is plausible that the tissue maintains its function for the most part even with low levels of Lox (~80% downregulation) and that more critical phenotypes emerge with further reduction of its expression or induction of hypertension. Altogether, linkage of both LOX extracellular and intracellular roles benefit our understanding of arterial development and pathologies.

## Materials and Methods

### Mice

All experiments involving mice conform to the relevant regulatory standards (AAALAC, Technion IACUC and national animal welfare laws, guidelines and policies). All mice are housed in IVC’s (Techniplast) according to space requirements defined by the NRC. All rooms are set to have 22 °C ± 2° and humidity of 30–70%. All HVAC parameters are controlled by a central computerized monitoring system. Light cycle is set to full light 10 h half-light 2 h and complete darkness 12 h, light cycle is monitored by the computerized central system. *Lox^fl^*mice were previously described ^15^. *SMMHC-CreER^T2^*(*MYH11:Cre^ERT2^*) ^14^ and *smMHC^Cre/eGFP^* (*MYH11:Cre*^eGFP^) ^25^ were purchased from JAX mice (JAX strains 019079 and 007742, respectively). Cre activation for *SMMHC-CreER^T2^*mice was carried out by injecting 50 µl tamoxifen (Sigma T5648, 20 mg/ml dissolved in corn oil) for 5 consecutive days to 3 week-old male mice, then 100 µl once a month. Hypertension induction was carried out via subcutaneous implantation of osmotic pumps (Alzet mini-osmotic pump Model 2004; 0.25µl/hour, 28 days) loaded with Angiotensin II (AngII; 10mg/1kg/day) to 3 month-old mice. Harvesting of aortas was performed 4 weeks after pump implantation.

### Cell Culture

HAOSMCs were purchased and cultured according to the supplier’s protocol (354-05A, Sigma-Aldrich). Cells were further grown to passage 6, and used for experiments up to passage 11. Smooth muscle cell growth media (311-500, Sigma-Aldrich) was used through all cell culturing. Kinase modulation: HAOSMCs were cultured and infected with shRNA (*shLOX* and *shCtrl*). Cells were trypsinized and seeded onto new dishes at 30% confluence. Two hours post seeding, the medium was replaced with fresh growth medium containing either Calyculin A (Alomone labs C-100, 1nM), MLCK inhibitor (Tocris 1885, 100 µM), Rho-Activator II (Cytoskeleton CN03, 2µg/ml), or ROCK inhibitor y-s7632 (Peprotech 1293823, 1 µM). Cells were washed and fixed with 4% paraformaldehyde after 24 hours (Rho-Activator, ROCK inhibitor), or 4 days (Calyculin A, MLCK inhibitor).

### Lentiviral Infections

HAOSMC infection, using viral particles generated in HEK293T cells, was carried out according to standard protocols as previously described ^34^. MISSION® shRNA lentiviral plasmids (Merck) were used for knockdown of *LOX* (*shLOX1* – TRCN0000045991, *shLOX2* - TRCN0000293898, *shLOX3* - TRCN0000286463) and a non-target shRNA was used for control (*shCtrl* – SHC016). Viral particles generated by the HEK293T cells were collected in smooth muscle cell growth media, and used to infect 1 million HAOSMC cells in a 10 cm dish. 24 hours post infection, the media was replaced and Puromycin selection was administered for the following 5-7 days.

### Histochemistry

Harvested mice aortas were fixed overnight in 4% paraformaldehyde, then embedded in paraffin blocks. 3 µm sections were stained with Masson’s Trichrome (MTC), Verhoeff’s Van Gieson (VVG), or Hematoxylin & Eosin.

### Immunohistochemistry

Section immunohistochemistry were performed according to standard protocols and as previously described ^35^. The following antibodies were used: anti-Tubulin (abcam ab4074, 1:300), anti-Smoothelin (ProteinTech 23567, 1:1000), anti-Transgelin (abcam ab14106, 1:1000), anti-Calponin (abcam ab46794, 1:500), anti-MyosinIIB (DSHB CMII23, 1:50), anti-Filamin A (abcam ab76289, 1:200). Staining for F-actin was done using eGFP-conjugated phalloidin (Life Technologies R37110, 1:1000).

### Western blot

Protein lysates harvested from mice aortas or HAOSMCs were subjected to SDS-PAGE analysis. Polyacrylamide gels were transferred to nitrocellulose membranes, which were then probed with the following antibodies: anti-LOX (Cell Signaling D8F2K, 1:1000), anti-Smoothelin (ProteinTech 23567, 1:1000), anti-Transgelin (abcam ab14106, 1:1000), anti-Calponin (abcam ab46794, 1:5000), anti-ROCK1 (abcam ab97592, 1:1000), anti-ROCK2 (abcam ab125025, 1:3000), anti-phospho-MLC2 (cell signaling 3674), and anti-MLC2 (cell signaling 3672). Loading was determined using an anti-P97 (1:5000, kindly provided by Ariel Stanhill, Open University, IL).

### Immunoprecipitation

Immunoprecipitations were done using 1 mg HAOSMC lysate, incubated with 2.5 µg anti-LOX (LS-C143168, LSBio) or IgG (011-000-003, Jackson Laboratories) as control, in a co-immunoprecipitation buffer (20 mM Tris-HCl [pH 7], 150 mM NaCl, 2 mM EDTA, 1% NP-40) O/N at 4°C. Pre-clearing was done by incubating 2.5 µg IgG with the lysate prior to incubation with the specific antibody. All antibodies were initially incubated with Dynabeads Protein G (1003D, Thermo Fisher Scientific) in PBS [pH 7.4], 0.02% Tween-20 for 1 hour to form the conjugated complex of beads with antibodies. After incubation with the lysate, protein complexes were washed three times in PBS and subsequently extracted with SDS loading buffer for 3 minutes at 95°C. All incubations were performed at 4°C with gentle rotation.

### RNA Isolation and cDNA Synthesis

HAOSMC pellets were incubated with TRizol reagent (92608, Ambion Life Technologies) to obtain cellular content. RNA was isolated using phase separation with Chloroform. The RNA was then precipitated with isopropanol and washed with 75% EtOH. Following RNA isolation, the RNA was reverse transcribed and cDNA was prepared using cDNA kit 5X All-In-One RT MasterMix (abm G490) according to the manufacturer’s instructions.

### Real-time Quantitative PCR

Real-time qPCR was performed with the following probes from Sigma Aldrich: LOX fw, CGGCGGAGGAAAACTGTCT; LOX rv, TCGGCTGGGTAAGAAATCTGA; LOXL1 fw, CCACTACGACCTACTGGATGC; LOXL1 rv, GTTGCCGAAGTCACAGGTG; LOXL2 fw, GGGTGGAGGTGTACTATGATGG; LOXL2 rv, CTTGCCGTAGGAGGAGCTG; LOXL3 fw, TGGAGTTCTATCGTGCCAATGA; LOXL3 rv, CCTGAGGCTTCGACTGTTGT; LOXL4 fw, CTGGGCACCACTAAGCTCC; LOXL4 rv, CTCCTGGATAGCAAAGTTGTCAT; GAPDH fw, CATGAGAAGTATGACAACAGCCT; GAPDH rv, AGTCCTTCCACGATACCAAAGT.

Power SYBER Green PCR Master Mix (Thermo Fisher Scientific 4367659) was used for all reactions. **Imaging.** Images were acquired using the Zeiss LSM 880 laser scanning confocal attached to Axio Examiner Z1 upright microscope (Zeiss Germany), with X63 NA1.4 oil immersion objective and lasers line 405nm, 488nm, 561nm, 633nm. The LSM 880 is controlled by ZEN Black 2.3 (Zeiss Germany). Images used for nuclei analysis were acquire with Z stack. Quantitative image analysis of the nuclei was performed by using Imaris software version 9.8.2 (Oxford Instruments, England). Each confocal image was 3D reconstructed and segmented. HAOSMC live cell imaging of was acquired using the Essen Bioscience Incucyte ZOOM, using X10 lens for 72 hours at 10 minute intervals.

### Transmission Electron Microscopy

shRNA infected cells were grown on aclar film pretreated with poly-L-lysine, washed and fixed for 1 hour with 2% Glutaraldehyde, 2% paraformaldehyde in 0.1 M Sodium Cacodylate buffer (pH 7.4) containing 3% sucrose. The cells were washed with Cacodylate buffer, post fixed for 30 min using 1% Osmium tetroxide in cacodylate buffer, washed with DDW and incubated in 1% Uranyl Acetate for 30 min. Following dehydration in graded ethanol series, the aclar films with cells were moved to fresh wells filled with Epon812 to start infiltration and embedding. After polymerization, aclar film was pealed from the block and transverse sections were prepared. For aorta imaging, the extracted tissue was cleaned, cut into ~2-3 mm thick rings, placed immediately in the fixation solution containing 2% Glutaraldehyde, 2% paraformaldehyde in 0.1 M Sodium Cacodylate buffer for 1 hour at RT and then moved to 4°c for 16 hours. The samples were washed, post fixed with 1% Osmium Tetroxide for 1 hour, washed and en block stained with 2% Uranyl Acetate for 60 min. Following dehydration using ethanol, the specimens were infiltrated with Epon812 for 3 days, oriented in silicon molds and after polymerization, crosswise sections were made.

Ultra-thin 75nm sections were cut using ultramicrotome UC7 (Leica), transferred to copper grids and viewed using Talos L120C Transmission Electron Microscope at accelerating voltage of 120 kV.

### Mass Spectrometry Proteolysis

from cell lysates-cell pellets were suspended in 9M Urea, 400mM Ammonium bicarbonate and 10mM DTT following three cycles of sonication. 20ug protein from each sample were reduced with 3mM DTT (60°C for 30 min), modified with 9mM iodoacetamide in 100mM ammonium bicarbonate (in the dark, room temperature for 30 min) and digested in 1.5M Urea, 66mM ammonium bicarbonate with modified trypsin (Promega) at a 1:50 enzyme-to-substrate ratio, overnight at 37oC. An additional second trypsinization was done for 4 hours. For IP: Protein G magnetic beads were suspended with 9M Urea, 400mM ABC, and incubated for 30’ at room temperature. Supernatant was transferred to a clean tube and reduced with 2.8mM DTT (60°C for 30 min), modified with 8.8mM iodoacetamide in 100mM ammonium bicarbonate (in the dark, room temperature for 30 min) and digested with Trypsin in 1.5M Urea, 66mM ammonium bicarbonate with 0.1 µg of modified trypsin (Promega) overnight at 37oC.

### Mass spectrometry analysis

The tryptic peptides were desalted using C18 tips (Top tip, Glygen) dried and re-suspended in 0.1% Formic acid. Peptides were then resolved by reverse-phase chromatography on 0.075 X 180-mm fused silica capillaries (J&W) packed with Reprosil reversed phase material (Dr Maisch GmbH, Germany). Peptides were eluted with different concentrations of Acetonitrile with 0.1% of formic acid: a linear 180 minutes gradient of 5 to 28% acetonitrile followed by a 15 minutes gradient of 28 to 95% and 25 minutes at 95% acetonitrile with 0.1% formic acid in water at flow rates of 0.15 μl/min. Mass spectrometry was performed by Q Executive HFX mass spectrometer (Thermo) in a positive mode (m/z 300–1800, resolution 120,000 for MS1 and 15,000 for MS2) using repetitively full MS scan followed by collision induces dissociation (HCD, at 27 normalized collision energy) of the 30 most dominant ions (>1 charges) selected from the first MS scan. The AGC settings were 3×106 for the full MS and 1×105 for the MS/MS scans. The intensity threshold for triggering MS/MS analysis was 1×104. A dynamic exclusion list was enabled with exclusion duration of 20 s. The mass spectrometry data was analyzed using the MaxQuant software 1.5.2.8 ^36^ for peak picking and identification using the Andromeda search engine, searching against the human proteome from the Uniprot database with mass tolerance of 6 ppm for the precursor masses and the fragment ions. Oxidation on methionine, Lysine and protein N-terminus acetylation were accepted as variable modifications and carbamidomethyl on cysteine was accepted as static modifications. Minimal peptide length was set to six amino acids and a maximum of two miscleavages was allowed. Peptide- and protein-level false discovery rates (FDRs) were filtered to 1% using the target-decoy strategy. Protein table were filtered to eliminate the identifications from the reverse database, and common contaminants and single peptide identifications. The data was quantified by label free analysis using the same software.

### Cell to ECM correlation

For brevity, cell to its surrounding ECM correlation was performed by processing the cell and its matrix separately. The cells having large and distinguished shape were segmented using a pre-trained convolutional neural network (CNN). In order to eliminate noise as much as possible, small aggregates were also segmented and omitted from further steps in the process.

The ECM was characterized with elongated but sometimes faint texture. Therefore, we commenced by enhancing high frequencies using Embos filter, followed by random walker segmentation which allows to strip out these fine structures. The ECM segmented structures were further refined according to their aspect ratio. Principal component analysis (PCA) helped define the angle of each piece of ECM. Using PCA, we similarly calculated the angle of cells, defining a new image of cells, where their gray level values represented their angle. Then, an envelope that follows cell’s outer shape was defined, leaving the cell and some safety margins out of this envelope. This envelope encompasses ECM structures, at which we calculated their median angle. At this point, we defined another cells image, where the gray level of each cell was the angle of its surrounding ECM. At the final stage, the correlation between cell’s angle and the angle of its surrounding ECM was calculated as the cosine of the angle difference between the cell and its surrounding ECM. Since both negative and positive values have the same meaning, the absolute value was taken.

### Live-cell Ca^2+^ imaging

HAOSMCs were seeded on poly-L-Lysine (PLL, 5µg/ml) coverslips 1 day prior to the experiment. Cells were loaded with 5µg/ml Fura-2 AM (Invitrogen), a fluorescent Ca^2+^ indicator, in Ca^2+^-free Ringer’s solution for 30 min at room temperature. Next, the cells were washed and transferred onto an imaging chamber and imaged using a fluorescence microscope. To determine the plasma membrane Ca2+ ATPase (PMLC) activity, the cells were treated with Ca2+-free Ringer’s solution containing 2μM ionomycin (Alomone labs) and 0.2mM EGTA. The rate of Ca2+ clearance from the cytoplasm was determined by applying the exponential decay function in R/Python software.

### p-MLC Image Analysis

Python and its image processing libraries were used to analyze segmented aortic images, focusing on quantifying red (p-MLC) regions. Using a (7, 7) kernel, we analyzed each image segment for mean red intensity. Histograms were generated to visualize intensity distributions, and the Kolmogorov-Smirnov test was used to compare the groups statistically.

### Mitochondrial Image Analysis and Segmentation

We utilized Python along with Detectron2 to conduct an in-depth analysis of images annotated for normal and abnormal mitochondria. Annotations were carried out using MakeSense.

## Author Contribution

R.A. conducted the majority of the experiments. S.Z.E. assisted in cell culture and western blot assays and A.O., A.S. and S.M. assisted in image analysis. A.K. assisted in aorta isolations and western blot analyses. R.K. and R.P. assisted in calcium-related assays. C.B.P. and S.K.G. assisted cell based assays and data analyses. R.A. and P.H planned the study, supervised and analyzed the data.

## Acknowledgement

The authors wish to thank the UM-Israel Partnership for Research to SKG and PH, Israeli Science Foundation (1072/13) and (1111/18), by the Rappaport Family Foundation to P.H and NHLBI R35HL161016 to SKG. We are grateful to M. Holdengreber from the BCF Bioimaging Center (Faculty of Medicine, Technion); to Lihi Shaulov for TEM imaging; to members of the animal facility for excellent technical assistance and to the Smoler Proteomics Center at the Technion.

## Supplementary Figures

**Supplementary Figure 1. SMC-specific Lox deletion following hypertension leads to aneurysm formation.** Western blot (A) and quantification (B) for Lox from aortas of control (*Lox^fl/fl^*) or Mutant (*Myh11:Cre^ERT2^; Lox^fl/fl^* or *Myh11:Cre^ERT2^; Lox^Δ/fl^*) with no AngII infusion.

**Supplementary Figure 2. *LOX* knockdown in HAOSMC affects multiple processes.** Western blot for LOX from HAOSMC lysates treated with LOX inhibitors (A). HAOSMCs treated with LOX inhibitors do not show a significant size difference. Immunostaining of *shCtrl* and *shLOX* HAOSMCs for Ki-67 (C). Red staining indicate Ki-67 positive cells. Quantification of Ki-67 positive cells (D). Live cell imaging markers of cell proliferation (E) and cell death (F) of *shCtrl* and *shLOX* HAOSMC show reduced proliferation and augmented cell death in *shLOX* cells. Confocal images of DAPI nuclei staining and their Imaris 3D models (G). Quantification of regular disc-like nuclei and amorphic nuclei shows an increase of the latter in *shLOX* cells (H).

**Supplementary Figure 3. HAOSMC secrete significant LOX and ECM.** Representative ECM protein fold changes between ECM secreted by *shCtrl* and *shLOX* cells (A). HAOSMCs cultured on plastic in high confluency elongate and form layers of cells with a distinct direction (B). Following removal of the cells, ECM directionality can still be observed (C). Black arrows indicate the ECM directionality. Western blot for LOX from HAOSMC conditioned media showing LOX is highly secreted from the cells (D). Three independent conditioned media are shown. Quantification of *shCtrl* and *shLOX* average cell area following their co-seeding with parental HAOSMCs (E).

**Supplementary Figure 4. LOX is physically associated with cytoskeletal proteins**. KEGG (A) and IPA (B) analyses of proteins immunoprecipitated with LOX and identified by LC-MS/MS show enrichment of cytoskeletal proteins. Cytoskeletal-associated clusters are highlighted.

**Supplementary Figure 5. Hypertension is required for inducing aneurysms following *Lox* deletion in SMC.** Male and female mice of the indicated genotypes following AngII infusion for 28 days develop multiple aneurysms throughout the aorta length.

**Supplementary Figure 6. Abnormal mitochondrial appearance in *Lox* mutant medial SMC.** Mitochondrial area (A), major axis (B) and eccentricity (C) were monitored in control *Myh11:Cre^EGFP^* and mutant *Myh11:Cre^EGFP^; Lox^fl/fl^* medial SMC. Significant differences were observed in the shape of the mitochondria but not their area between the two genotypes. n= 4 mice of each genotype.

