## supplemental Figures for "Coordination between cytoskeletal organization, cell contraction and extracellular matrix development, is depended on LOX for aneurysm prevention"

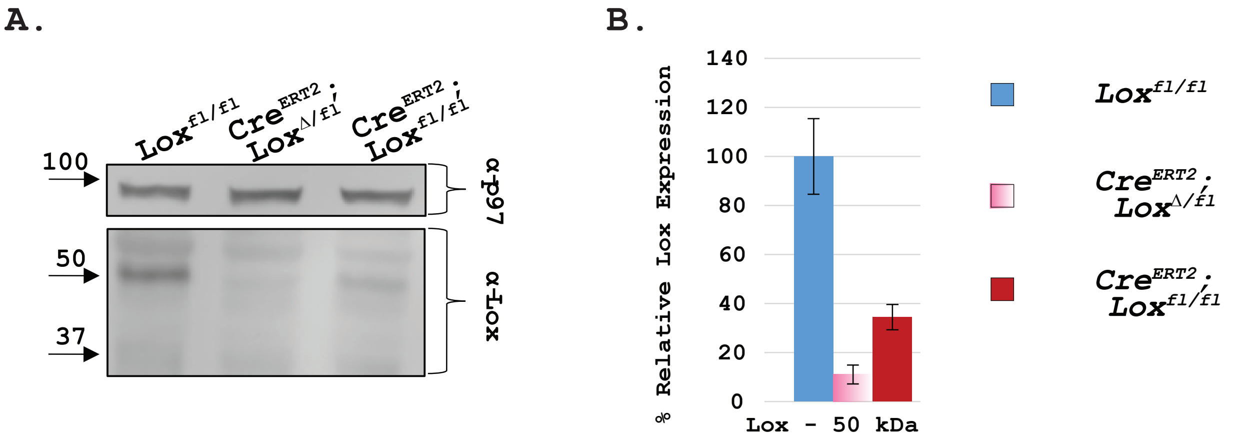


**Supplementary Figure 1. SMC-specific *Lox* deletion following hypertension leads to aneurysm formation.** Western blot (A) and quantification (B) for Lox from aortas of control (*Lox*$^{fl/fl}$) or mutant (*Myh11:Cre*$^{ERT2}$; *Lox*$^{fl/fl}$ or *Myh11:Cre*$^{ERT2}$; *Lox*$^{\Delta/fl}$) with no AngII infusion.

### Supplementary Figure 1

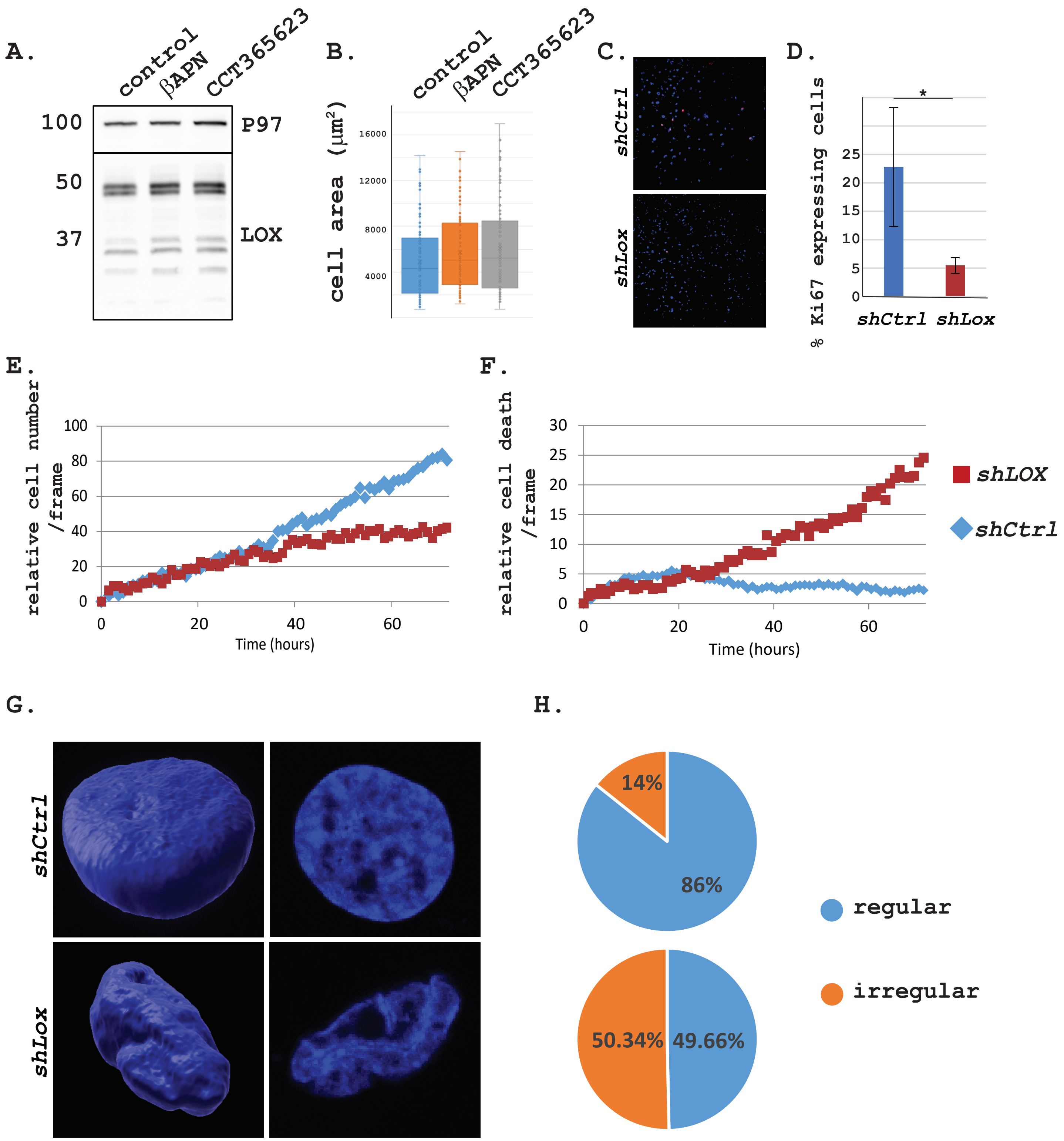


**Supplementary Figure 2. LOX knockdown in HAOSMC affects multiple processes.** Western blot for LOX from HAOSMC lysates treated with LOX inhibitors (A). HAOSMCs treated with LOX inhibitors do not show a significant size difference. Immunostaining of *shCtrl* and *shLOX* HAOSMCs for Ki-67 (C). Red staining indicate Ki-67 positive cells. Quantification of Ki-67 positive cells (D). Live cell imaging markers of cell proliferation (E) and cell death (F) of *shCtrl* and *shLOX* HAOSMC show reduced proliferation and augmented cell death in *shLOX* cells. Confocal images of DAPI nuclei staining and their Imaris 3D models (G). Quantification of regular disc-like nuclei and amorphic nuclei shows an increase of the latter in *shLOX* cells (H).

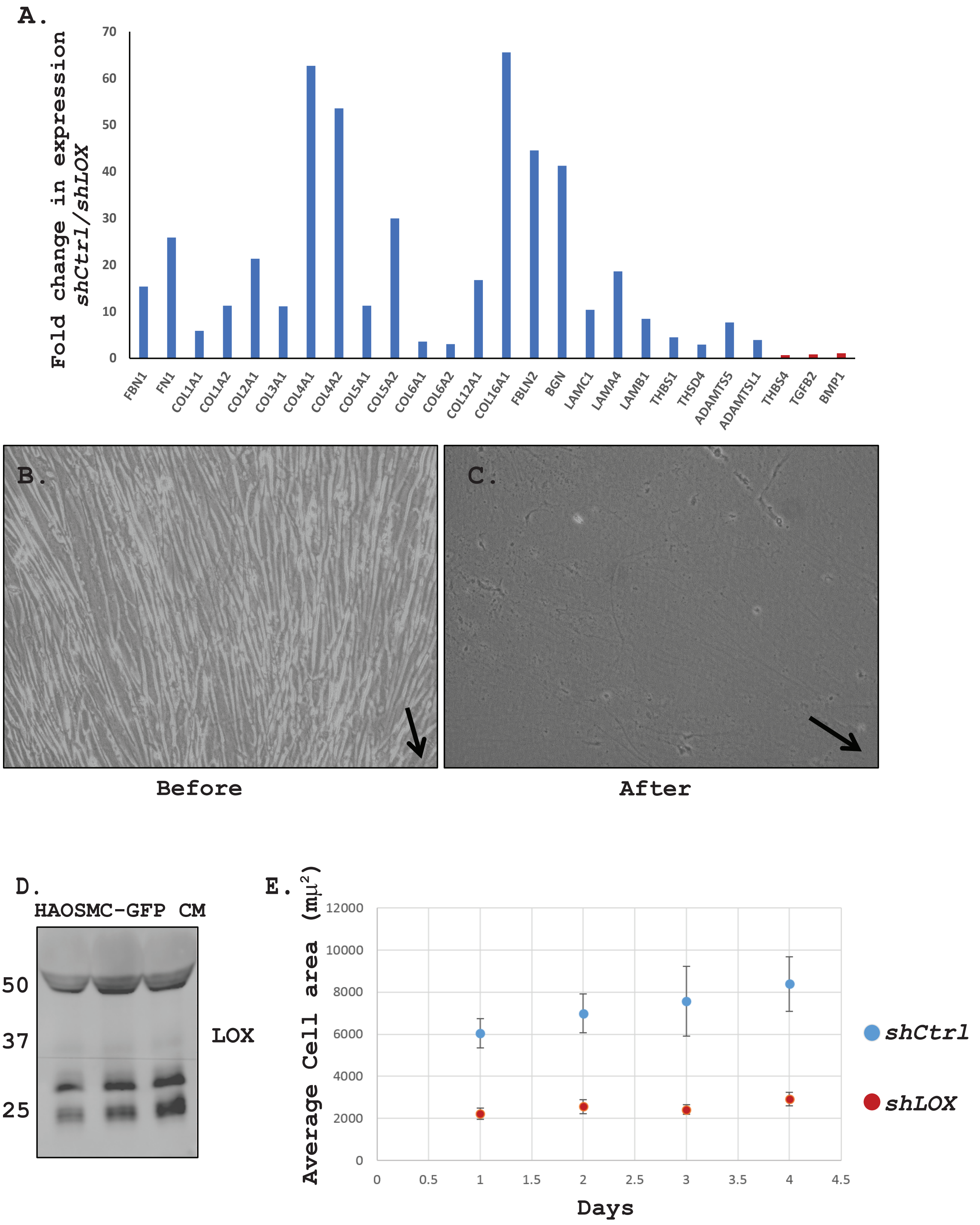


**Supplementary Figure 3. HAOSMC secrete significant LOX and ECM.** Representative ECM protein fold changes of ECM secreted by *shCtrl* and *shLOX* cells. Proteins in red are not significantly changed (A). HAOSMCs cultured on plastic in high confluency elongate and form layers of cells with a distinct direction (B). Following removal of the cells, ECM directionality can still be observed (C). Black arrows indicate the ECM directionality. Western blot for LOX from HAOSMC conditioned media showing LOX is highly secreted from the cells (D). Three independent conditioned media are shown. Quantification of *shCtrl* and *shLOX* average cell area following their co-seeding with parental HAOSMCs (E).

**Supplementary Figure 3**

A.

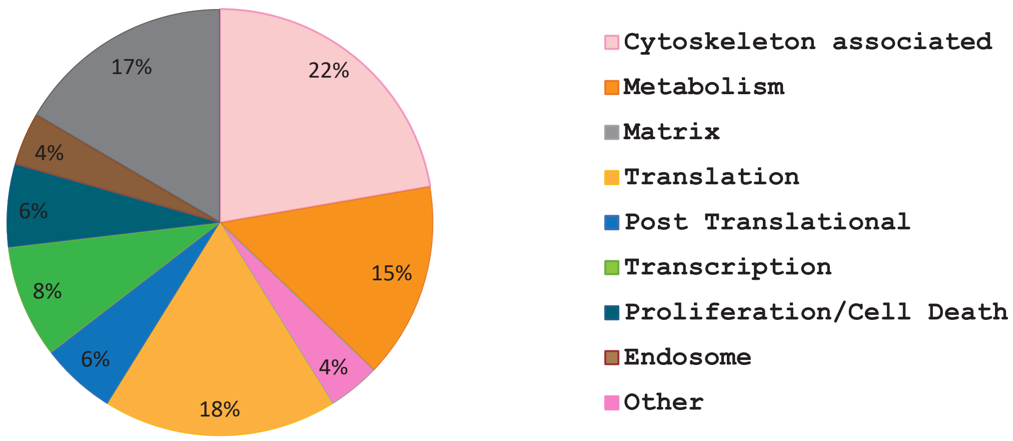


B.

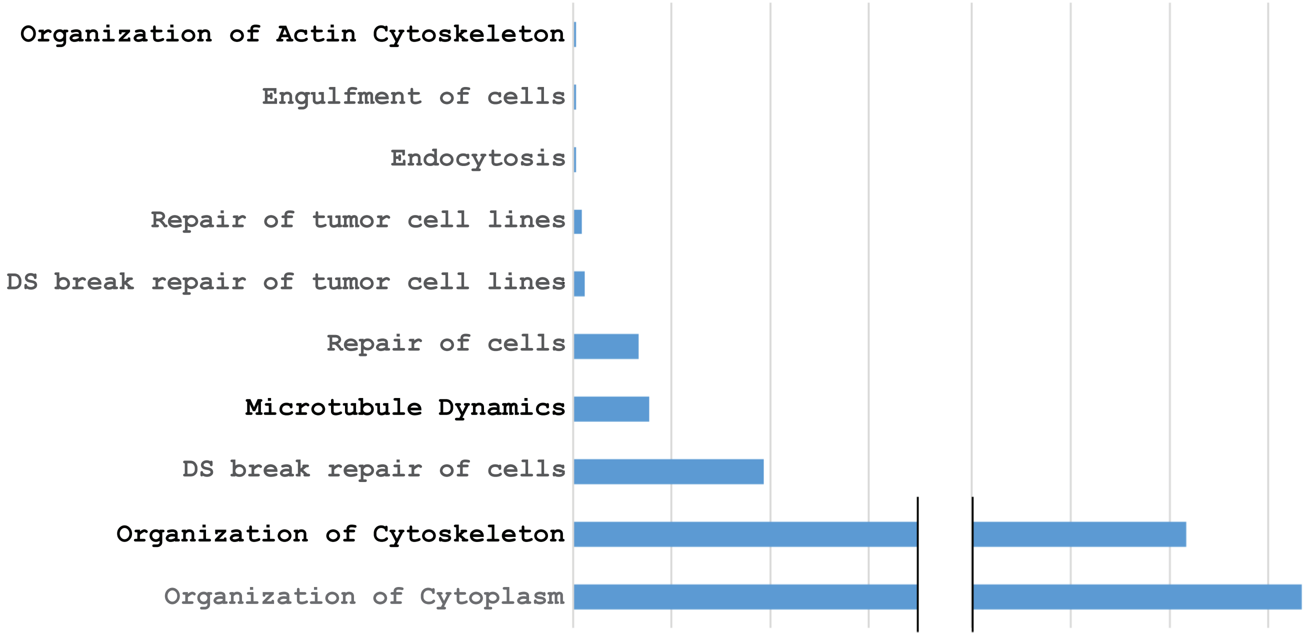


**Supplementary Figure 4. LOX is physically associated with cytoskeletal proteins.** KEGG (A) and IPA (B) analyses of proteins immunoprecipitated with LOX and identified by LC-MS/MS show enrichment of cytoskeletal proteins. Cytoskeletal-associated clusters are highlighted.

Supplementary Figure 4

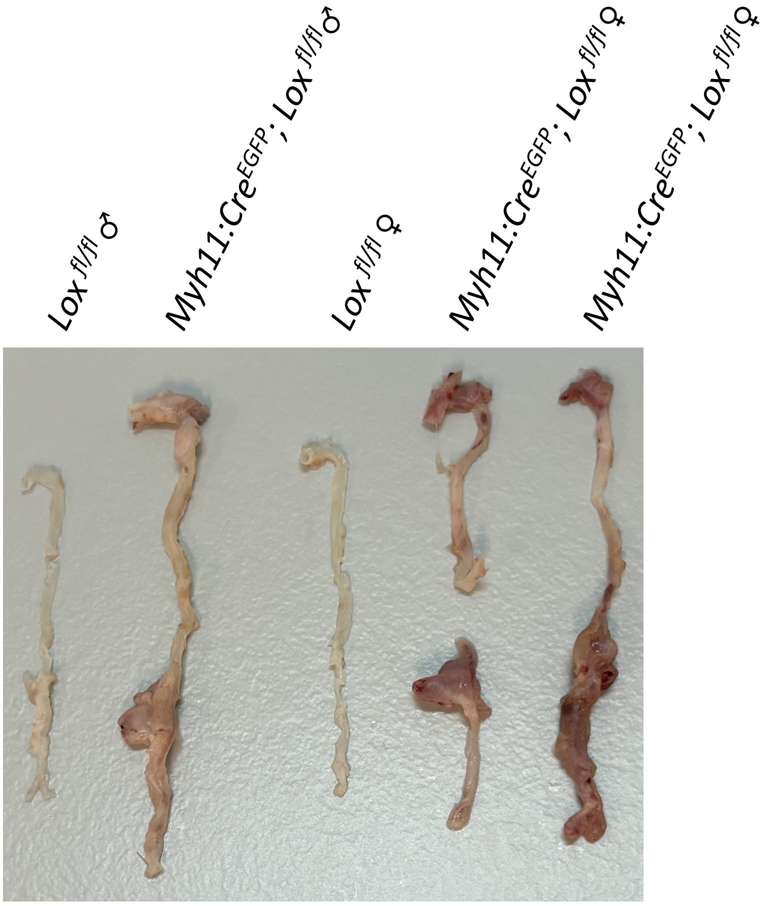


**Supplementary Figure 5. Hypertension is required for inducing aneurysms following *Lox* deletion in SMC.** Male and female mice of the indicated genotypes following AngII infusion for 28 days develop multiple aneurysms throughout the aorta length.

Supplementary Figure 5

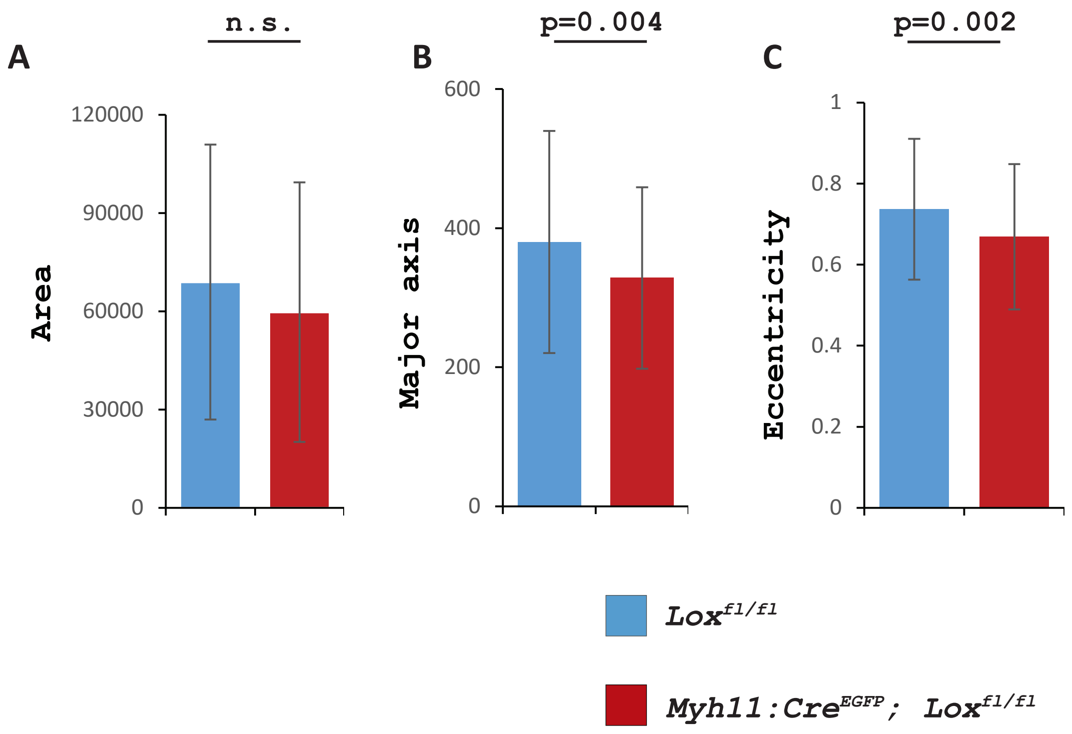


**Supplementary Figure 6. Abnormal mitochondrial appearance in *Lox* mutant medial SMC.** Mitochondrial area (A), major axis (B) and eccentricity (C) were monitored in control *Myh11:Cre*$^{EGFP}$ and mutant *Myh11:Cre*$^{EGFP}$*; Lox*$^{fl/fl}$ medial SMC. Significant differences were observed in the shape of the mitochondria but not their area between the two genotypes. n= 4 mice of each genotype.

Supplementary Figure 6
